# Environmental drivers of the decline of the fen orchid, Liparis loeselii

**DOI:** 10.1101/2024.02.22.581533

**Authors:** Dagmar Kappel Andersen, Rasmus Ejrnæs, Martine Minter, Tenna Riis, Erik Vinther, Hans Henrik Bruun

## Abstract

**Background:** *Liparis loeselii* is a rare and declining orchid species restricted to rich fens in the Northern Hemisphere. Habitat destruction, eutrophication, drainage and scrub encroachment have been suggested as reasons for the decline. However, which factors are most important is not well understood.

**Aims:** Based on vegetation and environmental properties from extant, potential and historical *L. loeselii* sites, we i) developed habitat suitability models from either Ellenberg Indicator Values or field-measured environmental properties, and ii) identified the primary reasons for the observed decline of *L. loeselii*.

**Results:** Nutrient status was the most important predictor for *L. loeselii* occurrence, followed by hydrology proxies (Ellenberg reaction and Ellenberg moisture). Vegetation height and Ellenberg light were of minor importance.

**Conclusion:** Effect partitioning based on sites, from which *L. loeselii* has gone locally extinct, pinpointed eutrophication and drainage to be the most likely primary drivers of the species’ demise. Phosphorus limitation induced by discharge of Calcium-rich groundwater seems to be crucial for *L. loeselii* to sustain populations in landscapes dominated by intensive agricultural.

## Introduction

Rich fens harbour exceptionally high plant species richness and constitute crucial habitat to many regionally rare and endangered species (Bedford and Godwin 2003; van Diggelen et al. 2006). Topographically, rich fens are found at positions in the landscape where nutrient poor water saturates the soil, e.g. in river valleys and dune slacks. The continuous supply of water creates a stable moisture regime throughout the year (Boeye and Verheyen 1992; Johansen et al. 2018), with relatively high pH and severe limitation of certain essential plant nutrients (Boomer and Bedford 2008; Wassen et al. 2005). The fen water is often groundwater, but may be nutrient poor, but base cation-rich surface water. Especially P limitation caused by high levels of Ca, Fe and Mg appears to plays a key role in maintaining rare and endangered species (Olde Venterink et al. 2001; Wassen et al. 2005).

The fen orchid, *Liparis loeselii* (L.) Rich., is a widespread Holarctic rich-fen species, but in its entire range rare and declining (Jacquemyn et al. 2023). The species is listed on Annex II and IV of the European Union Habitats Directive. As an Annex II species, *L. loeselii* requires designation of special protection areas within the Natura 2000 network, in addition to the union-wide strict protection enjoyed by Annex IV species (EC Habitats Directive 1992). Thus, according to the directive, EU member states are obliged to secure long-term “favourable conservation status” of *L. loeselii* populations. However, the habitat requirements of this species are not well understood. Therefore, management of occurrence sites is mainly based on “perceived best practice,” not on solid empirical inference.

In the EU member state Denmark, the number of historical *L. loeselii* populations was more than 100, from the occupancy has declined to 15 extant sites (Moeslund et al. 2023). The decline is commonly presumed to be caused by an overall decrease in the area and quality of suitable habitat as a consequence of drainage, eutrophication and abandonment of traditional management (Finderup Nielsen et al. 2021). However, there is little empirical knowledge on long term trends in the quantity and quality of potential *L. loeselii* habitat.

Past studies of *L. loeselii* have mainly focussed on its population biology (e.g. Jones 1998; McMaster 2001; Wheeler et al. 1998), including seed germination and mycorrhizal association (Illyés et al. 2005), and almost exclusively in dune slack habitats in Great Britain and the Netherlands. While studies of the population biology of the species are invaluable, they are not directly relatable to the distinction between suitable and unsuitable habitat. Also, conditions in dune slacks may differ in important environmental properties from inland rich fens. In general, population studies of *L. loeselii* have found declining population sizes, which in many cases has been attributed to natural succession in a dynamic dune ecosystem (Jones 1998; McMaster 2001). The study by Wheeler et al. (1998) of inland fen populations, suggested lowered water tables and lack of vegetation distrurbance, such as grazing, as drivers oif decline. To our knowledge, few studies have coupled *L. loeselii* occurrences with abiotic environmental properties to estimate habitat suityability and assess the reasons for the observed decline of the species (but see Wheeler et al. 1998; Grootjans et al. 2016 and Urban et al. 2020). Here, we investigate the occurrence of *L. loeselii* in response to habitat conditions related to soil fertility, moisture, pH and vegetation structure. Following from the above summarized studies, we *a priori* expect *L. loeselii* to be sensitive to increased levels of nutrients, in particular increased P availability, low light availability (caused by abandoned grazing and mowing), acidification and drainage.

The objectives of this study are to identify the main habitat constraints for *L. loeselii* based on a habitat suitability model and to evaluate current site management strategies. More specifically, the aim is to:

1. Develop a habitat suitability model for *L. loeselii* based on environmental indicators as well as field-measured habitat conditions.
2. Identify major threats to *L. loeselii* populations in Danish rich fens by evaluating data from historical (now locally extinct) and extant occurrence sites of *L. loeselii*.

## Methods

### Main data set

Our study area was Denmark. We included data from two sources: Rich-fen monitoring sites from The Danish National Monitoring Programme, NOVANA (https://novana.au.dk/), and supplementary data collection at occupied sites undertaken for the present study. We extracted data from all 285 sites in the monitoring programme 2004-2012, which included quadrats (plots) classified as fen in a broad sense (i.e. rich fen, alkaline spring, floating fen, wet meadow, humid dune slack and alkaline fen with *Cladium mariscus* (L.) Pohl. In a first step, we pruned the data so that for repeatedly monitored plots, only the latest recording remained, and monitoring sites with less than four survey plots were excluded. This procedure resulted in a data set of 17,436 vegetation quadrats. In a second step, individual plots of non-fen vegetation, which had been included due to the random placement of recording plots at monitoring stations, were excluded. This was done using a poisson regression of the presence of 27 typical rich fen species as a function of community-mean Ellenberg indicator values (Ellenberg et al. 1991) for moisture, nutrients and soil pH. Plots with effectively unsuitable habitat (too dry/wet, eutrophied or acidic plots) were defined as having a predicted number of typical fen species smaller than 0.5. No plots containing *L. loeselii* were excluded in this step.

Since monitoring plots are scattered randomly over sites and because *L. loeselii* is locally scarce, even within sites of occurrence, very few sampling quadrats included *L.loeselii* (i.e. 30 plots). In order to better cover the habitat variation of the focal species, we collected supplementary data at seven sites with *L.loeselii* populations (see Supplementary materials, Table S1). Five of the sites with supplementary plots were already part of the monitoring programme, both without *L. loeselii* in any sampled quadrats (Fig. 1). The supplementary data were collected in June through August 2013. After data trimming and addition of supplementary data, our data set contained 4,479 quadrats distributed over 270 sites of potentailly suitable habitat, of which 60 plots from 8 sites contained *L. loeselii*.

**Fig. 1.**
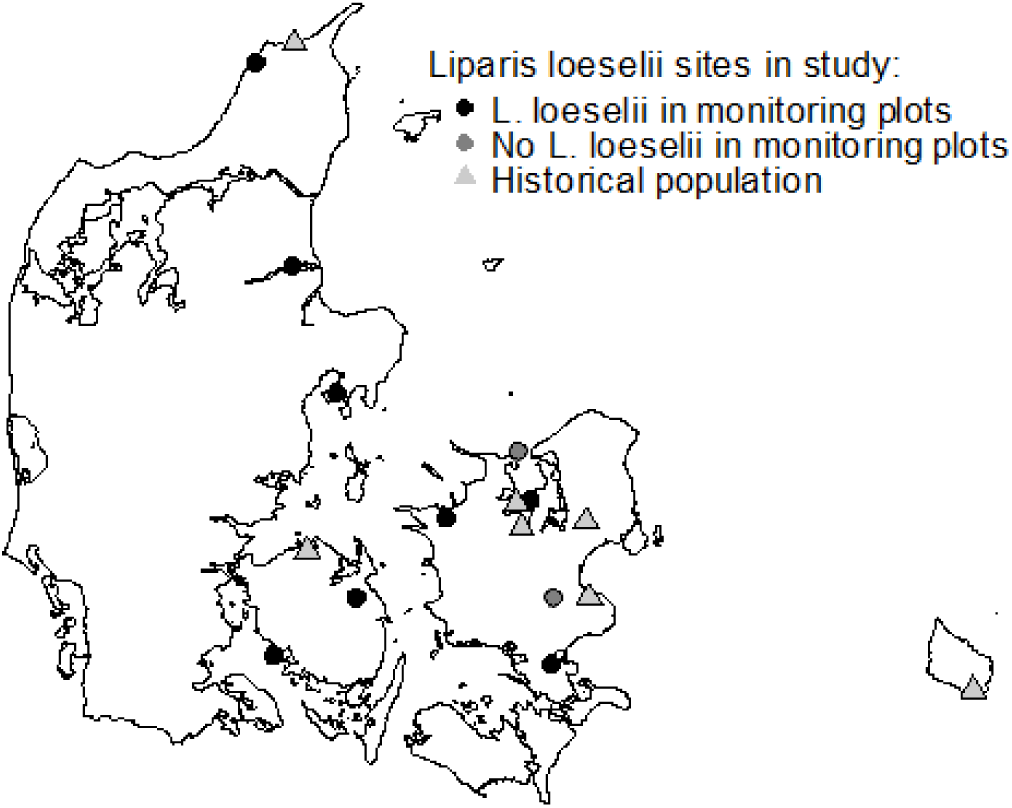
The 17 *Liparis loeselii* sites included in this study: 10 sites with *L. loeselii*, of which 8 sites had the species recorded in monitoring plots, and 2 sites only outside such plots. In addition, 7 sites harbouring historical, but now extinct, *L. loeselii* populations.

### Field survey and laboratory analyses

In the monitoring scheme, lists of vascular plant and bryophyte species were recorded along with visual assessment of vegetation structure from standard monitoring plots of 78.5 m^2^ (circle with radius = 5 m). Assessment of vegetation structure included vegetation height (calculated as an average of four measures in the center of the plot) and cover (in percentage) of trees and shrubs in two height classes (above and below 1 m tall). An identical protocol was used for the collection of supplementary data. GPS coordinates were available for all plots with corresponding data on elevation above sea level and a calculated Topographical Wetness Index, which is a proportional measure of the water retention potential in a given plot (Moeslund et al. 2013).

In a subset of plots, the following soil properties were assessed as part of the national monitoring scheme:

- pH was measured either – at wetter sites – directly in rhizosphere water using a combination glass electrode or – at drier sites – in a 0.01 M CaCl_2_ suspension of soil at 21°C with a combination glass electrode (available from 352 plots).
- Plant available soil P was measured (in 194 plots) by extracting fresh soil with 0.5 M NaHCO_3_, filtration, addition of H_2_SO_4_ and estimation of P concentration by spectrophotometry at 890 nm (following Banderis et al. 1976) (available from 194 plots).
- Nitrate in water was analysed with ion chromatography or flow injection analysis (available from 282 plots). The sample was injected in a stream of ammonium chloride and passed over a cadmium reactor, where nitrate was reduced to nitrite. Nitrite was reacted with sulphanilamide and naphtylethylenediamine and the resulting azo-colour was spectrophotometrically determined at 540 nm. Conductivity was measured in soil water in the field using a conductivity meter (available from 348 plots).
- Total N content in bryophyte tissue was measured using a LECO CNS-2000 analyser (Eurofins, Vejen, Denmark). Dried plant material was incinerated at 1100 °C. After measuring CO_2_, using infrared spectroscopy, N oxides were reduced to N using a copper catalyst, and CO_2_ was removed by absorption before measuring N in a thermal conductivity cell (available from 250 plots).
- Total P content in bryophyte tissue was measured using “Danish Standard” (DS259:2003/SM3120:2005-ICP-OES), in which plant material is treated with HNO_3_ and heated in a closed test tube at 120 °C and P content measured using atomic absorption spectrophotometry (Eurofins, Vejen, Denmark) (available from 201 plots).

*Calliergonella cuspidata* (Hedw.) Loesk. was collected as the standard species for tissue nutrient analysis but, except at a few sites (10 plots), from which *C. cuspidata* was absent and, therefore, *Campylium stellatum* (Hedw.) C. Jens. was collected instead. Based on NOVANA monitoring data and our own supplementary records, 201 plots included directly measured environmental variables, out of which 31 plots contained *L. loeselii*.

In addition to direct measurements, we used community-mean Ellenberg Indicator Values (Ellenberg et al. 1991) as indirect measures of soil pH (EIV-reaction), moisture (EIV-moisture), light availability (EIV-light) and nutrient availability (EIV-nutrient). In addition, we used the ratio EIV-nutrient/EIV-reaction (hereafter referred to as *nutrient ratio*), which takes the ubiquitous correlation between soil pH and soil fertility (e.g. Bragazza et al. 2002; Bridgham et al. 2001) into account and, thus, works as an indicator of eutrophication (Andersen et al. 2013).

### Data analysis

#### Habitat suitability models

Habitat modelling of *L. loeselii* was performed with generalized logistic regression as formulated by generalized additive modelling (GAM; R package mgcv, function: gam). GAM was preferred over a generalized mixed model, since GAM allows for non-linear relations, which were present in our data. Two sets of predictor variables were used: (i) directly measured environmental properties (Direct measures model, henceforth) and (ii) community-mean Ellenberg Indicator Values and vegetation structure (Indicator model, henceforth). The justification for an indicator-based model was that community-mean Ellenberg Indicator Values and vegetation height were available from all plots, whereas the directly measured variables were available for a subset of plots only. Smoothing was applied to all variables in order to allow for non-linear responses and in order to ensure comparable effect sizes between model terms. We initially used the default determination of optimal degree of smoothing (Wood 2006), but reduced the smoothing parameters further after examination of response curves aiming for biologically plausible responses reducing the risk of overfitting. We used cross-validation for variable selection (Burnham and Anderson 2002), in order to avoid overfitting due to pseudoreplication and spatial autocorrelation (Verbyla and Litvaitis 1989). In each case, the best model was chosen by leaving out all plots from one site at a time and evaluating model performance from the actual of *Liparis* incidence in the left-out plots. The best model was chosen based on the highest chi-square value and, furthermore, always had an even or higher number of correct predictions of *Liparis* occurrence compared to the second best model.

We used the best GAM model to predict the probability of *L. loeselii* in all plots. Variables excluded during model selection included: elevation, topographical wetness index, soil pH, nitrate in water, plant available P, and cover of trees and shrubs. For plant available P and nitrate in water, the weak explanatory power was probably related to very few measurements in plots with *L. loeselii* (3 plots).

In order to compare the goodness of fit of the Direct measures model and the Indicator model we used the subset of plots going into the Direct measures model and adjusted the probability criterion of predictions to reach the same the number of false and true positives (5 and 26). Hereby, we could compare the two models based on their proper discrimination between plots with observed absences.

#### Effect partitioning

In order to compare effect sizes of factors in the suitability model between different groups of plots (historical, extant and unsuitable sites), we extracted the effect of each predictor variable for each observation (plot). The effects from the logistic additive regression model cannot be readily interpreted before summation and back-transformation to probability scale, but they are centred on 0, which means that a positive partial effect values increase the probability of *L. loeselii* occurrence, whereas negative partial effect values decrease the probability.

#### Habitat constraints in historical sites

In the monitoring programme, seven sites are surveyed, from which *L. loeselii* have gone locally extinct since 1950. By comparing environmental conditions of these historical sites to sites with extant populations, we attempted to single out factors correlated with local extinction. From historical *L. loeselii* sites, all monitoring plots were included irrespective of present environmental conditions and occurrence of typical rich fen species. The Indicator model was used to predict the probability of *L. loeselii* occurrence.

## Results

### Habitat modelling

The Indicator model best predicting (after cross validation) *L. loeselii* occurrence included E-light, EIV-moisture, EIV-reaction and nutrient ratio (Chi-square value = 964.13, p < 0.0001). The effects of each explanatory variable on the probability of *L. loeselii* occurrence are shown in Fig. 2. Nutrient ratio, EIV-moisture and EIV-reaction were significant in the model, whereas EIV-light came out as marginally insignificant. Nutrient ratio had a linear negative correlation with *L. loeselii* occurrence probability, EIV-reaction had a positive correlation with saturation at high values whereas EIV-moisture and EIV-light had unimodal responses. *L. loeselii* probability peaked at EIV-light = 7.3 (semi-shade to full light), with lowest probability in shaded plots. The optimum EIV-moisture for *L. loeselii* was 7.9 (damp to wet) and lowest predicted probability is in the wettest plots (EIV-moisture=10, corresponding to shallow water). *L. loeselii* occurrence peaked at EIV-reaction = 6.3 (corresponding to neutral soil pH), levelling off at higher levels (Table 1, Fig. 2). All four factors had positive mean effects in plots with high probability (> 20%), but lower or negative mean effects in plots with low probability (< 1%) of *L. loeselii* occurrence, and especially mean effects of Nutrient ratio was markedly lower in plots with low probability (Fig. 4). A predicted probability of *L. loeselii* occurrence above 10% requires that the sum of factor effects in the logistic regression should exceed 7.19. All terms except nutrient ratio only contribute relatively small positive effect sizes, but in some cases contributed substantial negative values (Table 1, Fig. 2). Thus, in our Indicator model, a low nutrient ratio is essential to obtain a high probability of *L. loeselii* presence in a plot, but may not be sufficient if moisture, alkalinity or light conditions are suboptimal.

**Fig. 2.**
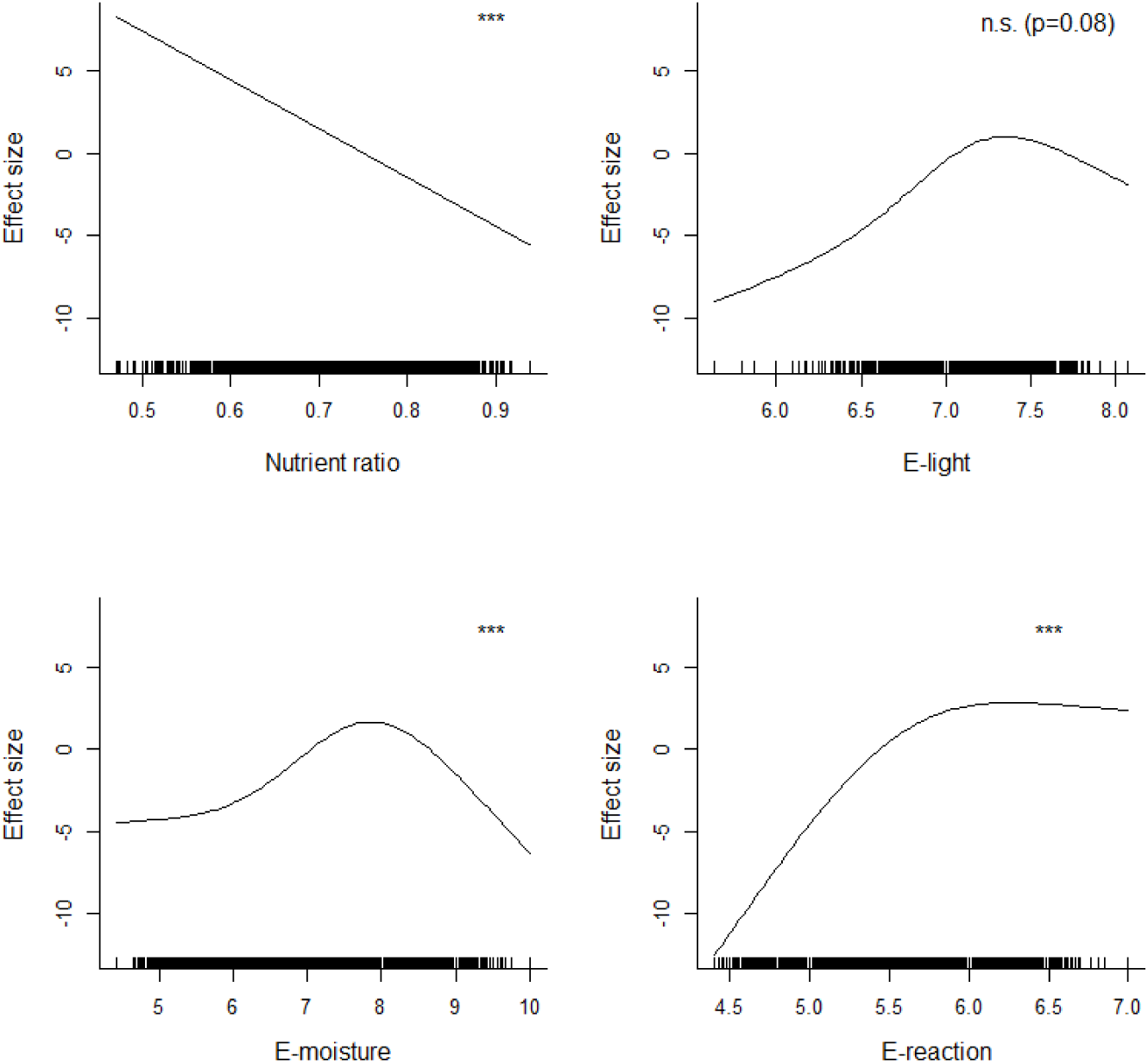
Effects of explanatory variables on the predicted probability of occurrence of *Liparis loeselii* in the Indicator model. Values above the horizontal zero line indicate higher probability of *L. loeselii* occurrence. Level of significance of each factors is indicated in the figure. ***: p-value < 0.001, **: p-value < 0.01, *: p-value < 0.05, n.s.= not significant. See methods for explanation of variable names.

**Table 1.** Summary of the maximum and minimum effect sizes, corresponding parameter value and linearity and estimated degrees of freedom (edf) of all factors in the Indicator model. The edf value reflects the degree of smoothing.

|  | Min. effect<br>(Parameter<br>value) | Max. effect<br>(Parameter<br>value) | P-value | Linearity<br>(edf) |
| --- | --- | --- | --- | --- |
| Nutrient ratio | -5.54 (0.94) | 8.30 (0.47) | <0.0001 | Linear (1) |
| EIV-light | -9.01 (5.64) | 1.02 (7.35) | 0.08 | Nonlinear (1.9) |
| EIV-moisture | -6.34 (10) | 1.65 (7.85) | 0.0001 | Nonlinear (2.7) |
| EIV-reaction | -12.5 (4.4) | 2.83 (6.27) | 0.0002 | Nonlinear (1.9) |

For the direct measures, the best model after cross validation included N concentration and N:P ratio in bryophyte tissue and vegetation height (Chi-square value = 39.90, p < 0.0001). All three factors were significant; however, vegetation height only marginally so. N concentration in bryophyte tissue was negatively correlated, whereas N:P ratio in bryophyte tissue was positively correlated with *L. loeselii* probability, indicating that low nutrient status and especially P availability is essential to the presence of *L. loeselii* (Fig. 3). We found no significant difference between N:P ratio measured in the tissue of *C. cuspidata* and *C. stellatum*. Vegetation height had a nonlinear response with the lowest probability at a vegetation height around 50 cm (Table 2, Fig. 3). All three factors had positive mean effects in plots with high probability (> 20%), but negative mean effects in plots with low probability (< 1%) of *L. loeselii* occurrence (Fig. 5). In particular, N:P ratio was much higher in plots with high occurrence probability.

**Fig. 3.**
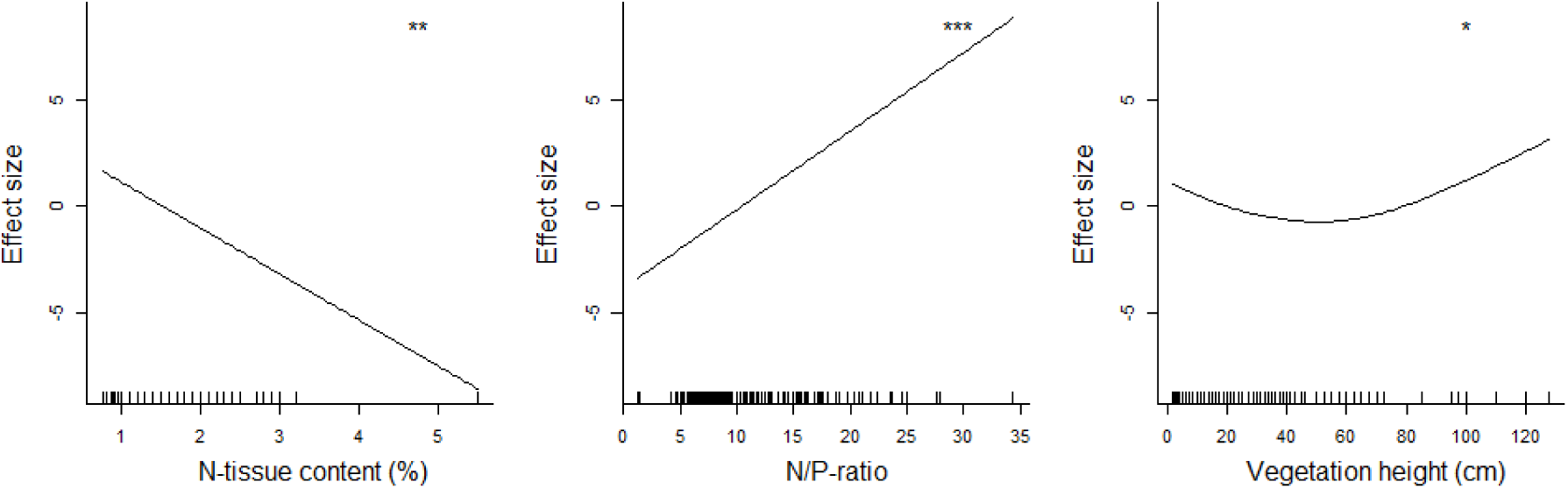
Effects of explanatory variables on the predicted probability of occurrence of *Liparis loeselii* in the Direct measures model. Values above the horizontal zero line indicate higher probability of *L. loeselii* occurrence. Level of significance of each factor is indicated in the figure. ***: p-value < 0.001, **: p-value < 0.01, *: p-value < 0.05. See methods for explanation of variable names.

**Fig. 4.**
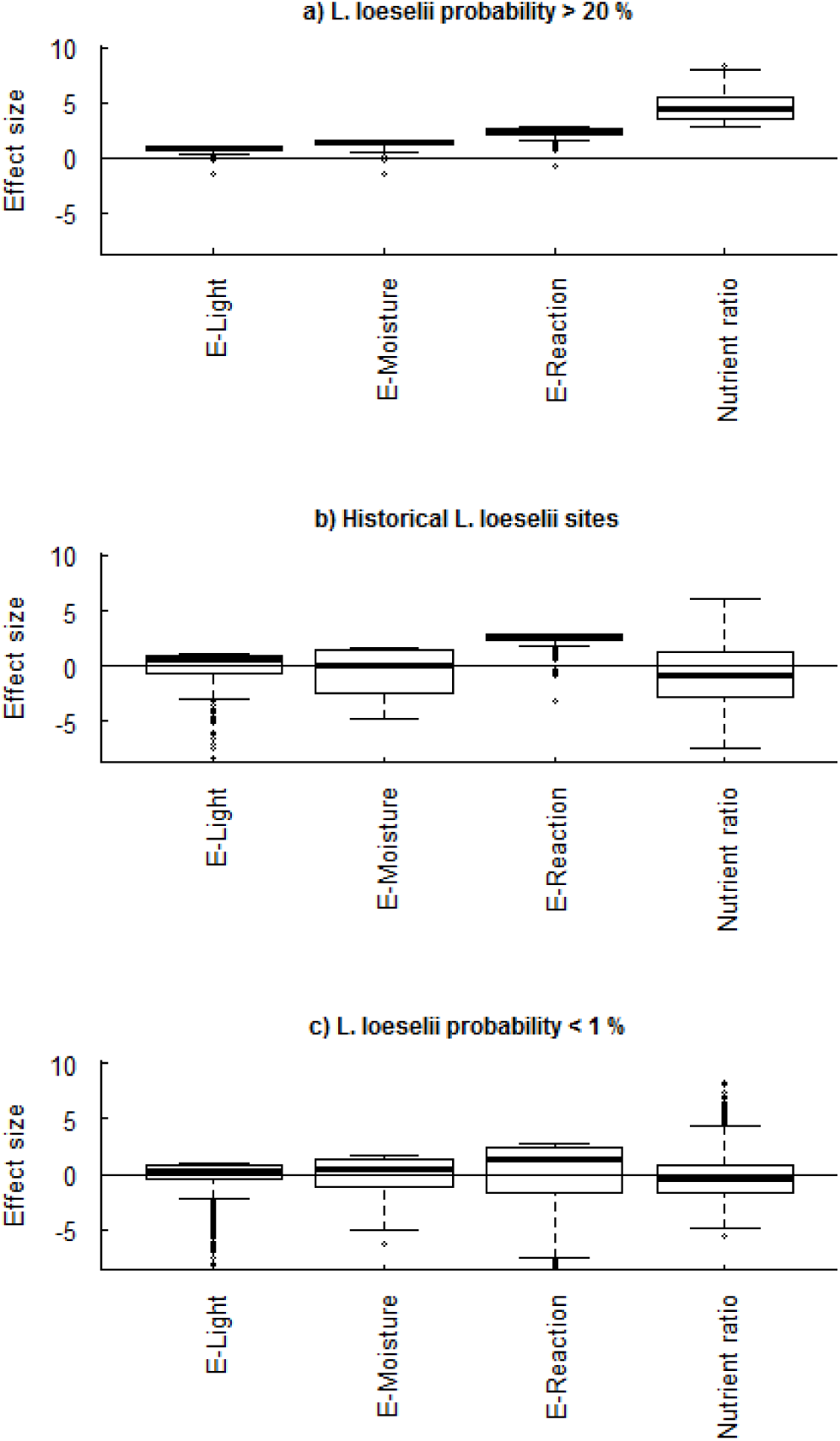
Effect sizes of model factors in plots with with a predicted probability of *Liparis loeselii* occurence > 20 % (a), historical sites of *L. loeselii* (b) and plots with a predicted probability of *L. loeselii* occurrence < 1% (c). Positive values increase the probability of *L. loeselii*, whereas negative values decrease the probability.

**Fig. 5.**
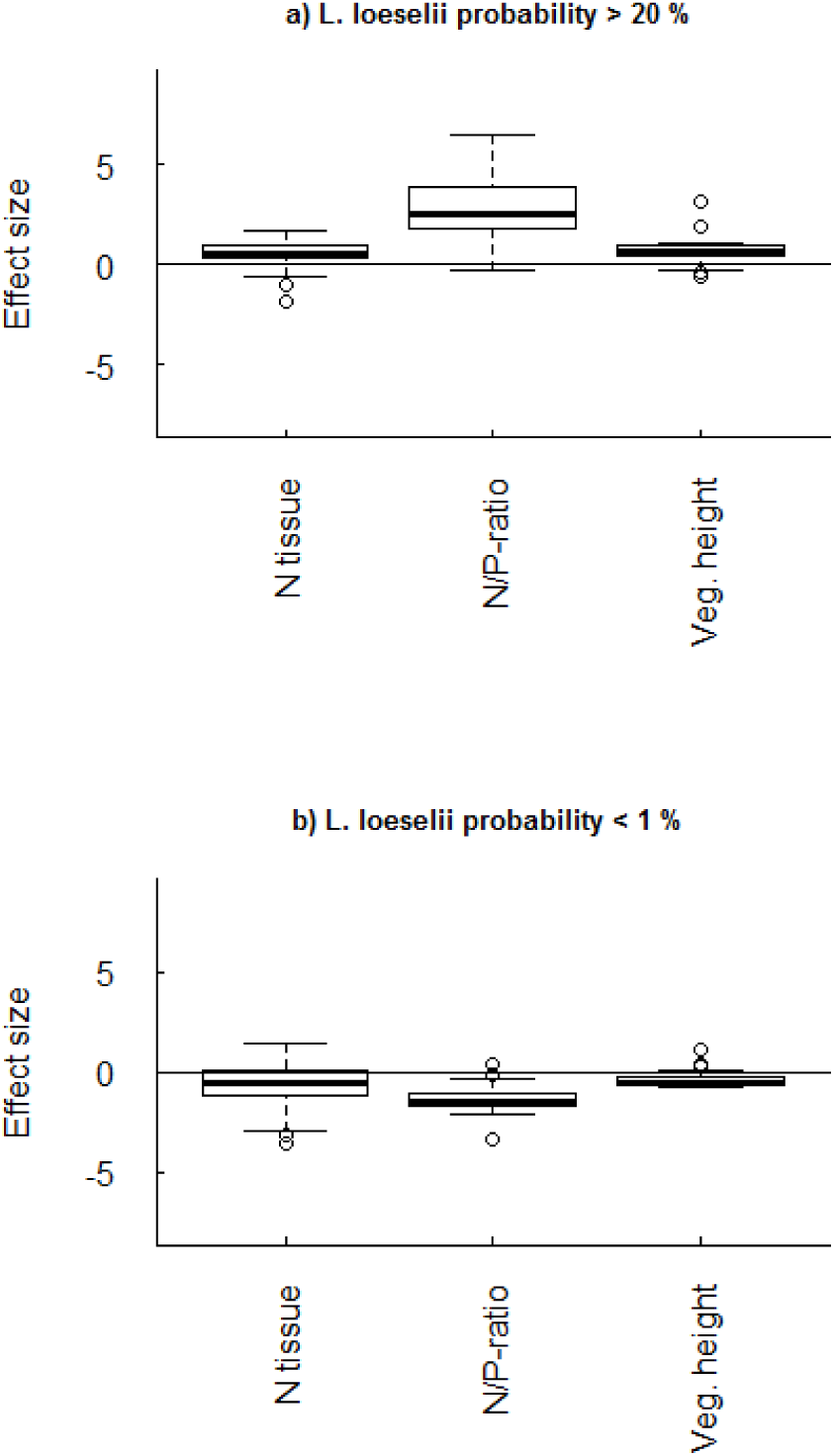
Effect sizes of model factors in the Direct measures model for plots with *Liparis loeselii* probability over 20% (a) and below 1% (b). Positive values enhance the probability of *L. loeselii*, whereas negative values decrease the probability.

**Table 2.** Summary of the maximum and minimum effect sizes, corresponding parameter value and linearity and estimated degrees of freedom (edf) of all factors in the Direct measures model. The edf value reflects the degree of smoothing.

|  | Min. effect<br>(Min. value) | Max. effect<br>(Max. value) | P-value | Linearity<br>(edf) |
| --- | --- | --- | --- | --- |
| N tissue concentration | -8.59 (5.5) | 1.66 (0.76) | 0.003 | Linear (1) |
| N:P ratio | -3.38 (1.24) | 8.86 (34.38) | <0.0001 | Linear (1) |
| Vegetation height | -0.73 (52.5) | 3.12 (127.5) | 0.047 | Nonlinear (2.2) |

Comparing the two different models after locking of the performance with regard to predictions of observed occurrences, revealed that the Direct measures model resulted in an appreciably higher number of false positives than the Indicator model (Table 3). The correlation between the predictions of the two models was relatively high (Spearman rank correlation coefficient = 0.76, p < 0.0001) and also between the two strongest variables, namely nutrient ratio and N:P ratio in tissue (Spearman rank correlation coefficient = -0.61, p < 0.0001), indicating that the Indicator model based on community-mean Ellenberg values reflects real conditions (Direct measures model).

**Table 3.** The observed and predicted numbers of *Liparis loeselii* presences when 5 false negatives are accepted. The cut value of predicted probability corresponding to 5 false negatives in each model is given.

|  | Observed/<br>predicted | Indicator<br>model | Direct measures<br>model |
| --- | --- | --- | --- |
| True negative | 0/0 | 162 | 111 |
| False positive | 0/1 | 8 | 59 |
| False negative | 1/0 | 5 | 5 |
| True positive | 1/1 | 26 | 26 |
|  | Cut value | 10% | 5.9% |

### Historical sites

Comparing the modelled effects of factors in the seven historical, but now unoccuopied, *L. loeselii* sites to extant sites, the major differences were the nutrient ratio and EIV-moisture (Fig. 4). The nutrient ratio eutrophication indicator was markedly higher at historical sites and the negative effect of EIV-moisture in historical plots arose mainly from plots being too dry (118 plots were drier than optimum, whereas eight plots were wetter). EIV-reaction, vegetation height and EIV-light had mean values more or less in the range of plots with extant *L. loeselii* (Fig. 4). The tendencies in effects at historical sites were very similar to the effects in NOVANA-plots with low occurrence probability of *L. loeselii* (Fig. 4).

## Discussion

We found that infertile conditions – probably P limitation in particular – is essential to the occurrence of *L. loeselii*. However, additional important predictors were soil pH and soil moisture, and to a lesser degree vegetation structure and light availability. Further, we found congruence between models based on directly measured variables and indicator models based on community-mean Ellenberg Indicator Values. Our models suggested that local extinction of *L. loeselii* in Denmark in recent decades has been caused primarily by eutrophication and drainage.

### Habitat modelling – Important factors

The expectations of nutrient status, hydrology and vegetation structure as being important for *L. loeselii* were supported by our results. Environmental properties related to nutrient status (Nutrient ratio and directly measured N:P ratio) were the most important in predicting the occurrence of *L. loeselii*. To our knowledge, the relationship between *L. loeselii* occurrence and nutrient status has not been investigated directly elsewhere. However, a phytometric assay in an area with *L. loeselii* populations found the substratum of *L. loeselii* to be relatively infertile (Wheeler et al. 1992). Furthermore, several studies have underpinned the importance of low nutrient availability to rich fen plant diversity and occurrence of rare habitat specialist species (e.g. Bedford et al. 1999; Wassen et al. 2005). Moreover, infertile wetlands appear in general to support rare species much more often than fertile sites (Moore et al. 1989; Wheeler and Shaw 1991), especially where P is the main limiting factor (Olde Venterink et al. 2001, 2003; Wassen et al. 2005). In Denmark and elsewhere, extraction of groundwater and drainage has diminished the influence of Ca- and Fe-rich groundwater by reducing the discharge and the retention time, respectively. This has resulted in radical changes to moist-wet P limited habitats, and many plant species adapted to such conditions are now endangered. Several studies have used N:P ratio of aboveground biomass as a measure of nutrient limitation in freshwater wetlands (Bedford et al. 1999; Güsewell et al. 2003; Koerselman and Meuleman 1996; Wassen et al. 2005). Since the correlation between N:P ratio in the particular bryophyte species used here and ecosystem N:P limitation is unknown, we cannot compare our findings directly to the Redfield ratio (N:P ratios above 16 as P limited, below 14 as N limited and between 14-16 as co-limited; Koerselman and Meuleman 1996). Güsewell et al. (2003), however, found that P limited stands of bryophytes in Dutch fens and dune slacks had N:P ratios higher than 14, suggesting that N/P values of bryophytes are comparable to vascular plants. Hence, our results clearly indicate that a high N:P ratio in tissue obtained by low N concentration and very low P concentration is vital for *L. loeselii*, and this result definitely points to P limitation as a driving factor for *L. loeselii* occurrence. The seemingly low importance of measured plant-available soil P in this study is probably a consequence of the rather few available measurements from plots with *L. loeselii*, which is a byproduct of random plot placement in the national monitoring program.

Our results also suggest that *L. loeselii* is dependent on a stable groundwater discharge. EIV-reaction and EIV-moisture constitute two indirect estimates of hydrology in the models because high EIV-reaction in fens indicates a high pH, which is mainly achieved by supply of Ca-rich groundwater. The effect of increasing EIV-reaction on *L. loeselii* incidence is positive, pointing to the importance of high Ca content and thereby low soil P availability. P limitation is often maintained by chemical adsorption of P in Ca or Fe rich soils (Grootjans and van Diggelen 1995). Reduced groundwater flow has been found to result in lowered Ca and Fe supply, causing a shift from P limitation to N limitation (Olde Venterink et al. 2003). Our modelled probability of *L. loeselii* occurrence appears to peak at an EIV-moisture index value around eight, which corresponds verbatim to somewhere between “constantly moist, but not wet” and “water-saturated badly aerated soils” (Ellenberg et al. 1992). It seems plausible that many rare species are adapted to P limited conditions and are unable to compete when nutrient limitation is shifted to N limitation, which may be the consequence of an altered groundwater regime.

Contrary to expectations, no significant relationship between *L. loeselii* occurrence and two estimates of light availability was found. However, the indicator-based EIV-light contributed suficient explanatory power to be included in the overall model. The general focus of site management by grazing or mowing suggest a positive effect of light availability on *L. loeselii* occurrence probability. To short-stature herbs like *L. loeselii*, light availability is likely linked to vegetation height, which again is positively related to above-ground biomass (Axmanová et al. 2012). Further, several studies have shown threatened species to generally occur at relatively low-productive sites with low standing biomass (Moore et al. 1989; Olde Venterink et al. 2003; Wheeler and Shaw 1991). The lack of a clear effect of light in our habitat suitability model puts a question mark to the conventional reasoning outlined. We propose that low P availability may be a more decisive factor than vegetation height to *L. loeselii*. This is in line with the finding by Olde Venterink et al. (2001) that the density of threatened species increased with decreasing P availability regardless of whether P availability is correlated with vegetation height, which may be manipulated by grazing and mowing. This aspect deserves more attention from scientists and conservation managers alike.

We found a high degree of congruence between the Indicator model and the Direct measures model. Ellenberg indicator values as an indirect measure of ecological factors are used in numerous studies (e.g. Axmanová et al. 2012; Diekmann 2003; Hill and Carey 1997), and the reliability of using these surrogate variables have been discussed in several studies (e.g. Diekmann 1995; Schaffers and Sýkora 2000; Wamelink et al. 2002; Dwyer et al. 2021). It has been shown that Ellenberg values of nutrient status, soil reaction and nutrient ratio are valuable indicators when assessing conservation status in groundwater dependent terrestrial habitats (Andersen et al. 2013). Direct environmental measures are in principle preferred, because Ellenberg indicator values are derived from the species data to be modelled or interpreted, and may potentially lead to a circular reasoning. However, since indicator values are extracted from simple species lists, such data are more readily available than direvct measurements. Also, direct measures, e.g. tissue nutrient content, are often costly to obtain at sufficient accuracy (Andersen et al. 2013; Hájek and Hekera 2004). The high degree of congruence between the Indicator model and the Direct measures model in the present study supports the use of Ellenberg indicator values as a cheap and reliable approximation of real environmental conditions when only vegetation data is available.

Nutrient ratio (EIV-nutrient/EIV-reaction) has previously been shown to be a strong indicator of eutrophication, and thus for the conservation status, of fens and alkaline springs (Andersen et al. 2013). In accordance with the mentioned study, we found nutrient ratio to be a good predictor of the presence of *L. loeselii*. The rationale for using *nutrient ratio* rather than EIV-nutrient alone is that EIV-nutrient correlates strongly with EIV-reaction because fertile places in European ecosystems were always relatively base-rich and alkaline. By correcting for this correlation, it is possible to properly detect infertile but also alkaline habitats. . A low nutrient ratio thus indicates that the average species present shows preference for low nutrient status relative to its pH preference. The strong negative correlation found in our study with N:P ratio (Spearman rank correlation coefficient = -0.61) was expected, since the P-tissue concentration is to a large extend controlled by Ca concentration in the soil and soil water. Thus, for rich fens, *nutrient ratio* seems to work as a reliable proxy variable for N:P ratio, when only species composition data are available.

### Local extinction of Liparis in Danish rich fens

Effect partitioning of environmental properties in historical *L. loeselii* sites assign eutrophication and drainage as the primary reasons for the demise of *L. loeselii*. Unlike most orchids, *L. loeselii* is generally viewed as a pioneer plant tracking recent disturbances in dynamic ecosystems (Grootjans et al. 2017). Also, low genetic differentiation between geographically distant Danish *L. loeselii* populations suggests good dispersal and colonization potential (Andersen et al. 2005). Hence, dispersal limitation is unlikely to be a major constraint to *L. loeselii* incidence. Consequently, lack of suitable habitat seems a more plausible explanation for the rarity of the species. At the seven historical *L. loeselii* sites investigated here, high nutrient ratio and dry site conditions were the primary explanations for low habitat suitability. Drainage and eutrophication may, however, be closely connected, since aeration of peat increases N and P mineralization and availability (Mettrop et al. 2014).

### Perspectives of grazing and mowing as management tools in extant and potential *L. loeselii sites*

Our study suggests that vegetation height is of secondary importance to the occurrence of *L. loeselii* compared to nutrient availability and hydrology. Throughout Central and Northern Europe, grazing and mowing are mainstream conservation management tools, employed to maintain or restore species richness in grasslands, including fens. It is often held that the mechanism goes via nutrient removal (Kooijman and Smit 2001). However, and in agreement with our findings, several studies have stressed that – despite a positive effect of mowing on species richness – the effect is minor compared to the effect of hydrology (Fojt and Harding 1995; van Belle et al. 2006; van Diggelen et al. 1996; Ilomets et al. 2010). Fojt and Harding (1995) concluded that “changes in hydrology cannot be compensated for by increased management”. Similarly, van Loon et al. (2011) found that the most efficient strategy to counteract fen deterioration is to eliminate drainage ditches in order to allow the horizontal movement of groundwater through the peat layer. Management by grazing and mowing is unlikely to restore an infertile environment, at least in the short-medium term, but may restore optimal light conditions after hydrological conditions have restored suitable nutrient status. Although the establishment and survival of *L. loeseli* is likely to be favoured by moderate levels of vegetation disturbance by grazing or mowing, this effect cannot justify artificial drainage in order to secure livestock grazing.

## Conclusion

The decline of *L. loeselii* in rich fens in Denmark was found to be primarily explained by eutrophication and drainage. Thus, infertile conditions – Phosphorus limitation in particular – is crucial to maintain or restore rich fens with threatened and rare species, such as *L. loeselii.* Natural hydrology, including the the impact of Calcium-rich groundwater, seems to be of particular importance and deserves priority over management by grazing or mowing.

## Supporting information

Table S1. Sites from which supplementary sample quadrats including Liparis loeselii were obtained

## Acknowledgements

We thank Jesper Moeslund for helping with database extraction, Peter Wind for sharing knowledge of populations and Annita Svendsen, Anders Juel and Johanne Fagerlind for assisting in the field.

## Author contributions

DKA - Conceptualization, Methodology, Data curation, Investigation, Formal Analysis, Visualization, Writing – original draft; RE - Conceptualization, Methodology, Investigation, Formal Analysis, Funding acquisition, Supervision, Writing – review & editing; TE - Supervision, Writing – review & editing; EV - Methodology, Investigation, Funding acquisition, Writing – review & editing; HHB - Conceptualization, Methodology, Investigation, Writing – original draft.

