## Supplementary material for "Environmental drivers of the decline of the fen orchid, Liparis loeselii": Table S1. Sites from which supplementary sample quadrats including Liparis loeselii were obtained

**Table S1**. Sites from which supplementary sample quadrats including *Liparis loeselii* were obtained in the summer of 2013.

| Site name | Geolocation | NOVANA station | Number of quadrats |
| --- | --- | --- | --- |
| Vandplasken | 57.518°N 9.880°E | yes | 4 |
| Nørlev Kær | 57.514°N 9.867°E | yes | 2 |
| Ajstrup Kær | 56.706°N 10.142°E | yes | 4 |
| Tved Kær | 56.199°N 10.453°E |  | 4 |
| Urup Dam | 55.385°N 10.592°E | yes | 4 |
| Helnæs Made | 55.147°N 10.008°E | yes | 4 |
| Even | 55.121°N 11.996°E |  | 4 |
| Forklædet | 55.774°N 11.841°E | yes | 5 |

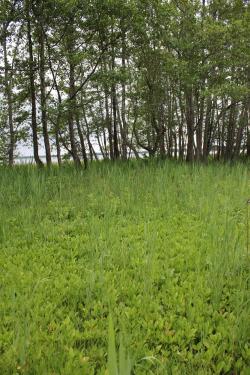

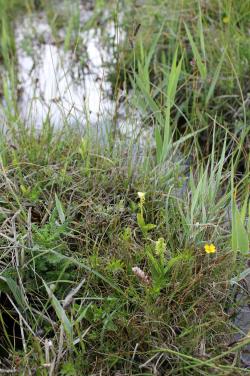

Fig. S1. Habitat of *L. loeselii*. To the left, Vandplasken, where the species co-occur with *Herminium monorchis*, *Potentiella erecta*, *Phragmites australis* and many other species on rich fen tussocks. To the right, Ajstrup Kær, in which *L. loeselli* occurs in moss-rich vegetation with visual dominance of *Menyanthes trifoliata*. Photos: Dagmar Kappel Andersen.
